# Genetic basis of variation in heat and ethanol tolerance in *Saccharomyces cerevisiae*

**DOI:** 10.1101/318212

**Authors:** Linda Riles, Justin C. Fay

**Affiliations:** Department of Genetics, Washington University, St. Louis, MO 63110

**Keywords:** quantitative trait, mapping, yeast, natural variation

## Abstract

*Saccharomyces cerevisiae* has the capability of fermenting sugar to produce concentrations of ethanol that are toxic to most organisms. Other *Saccharomyces* species also have a strong fermentative capacity, but some are specialized to low temperatures, whereas *S. cerevisiae* is the most thermotolerant. Although *S. cerevisiae* has been extensively used to study the genetic basis of ethanol tolerance, much less is known about temperature dependent ethanol tolerance. In this study, we examined the genetic basis of ethanol tolerance at high temperature among strains of *S. cerevisiae*. We identified two amino acid polymorphisms in *SEC24* that cause strong sensitivity to ethanol at high temperature and more limited sensitivity to temperature in the absence of ethanol. We also identified a single amino acid polymorphism in *PSD1* that causes sensitivity to high temperature in a strain dependent fashion. The genes we identified provide further insight into genetic variation in ethanol and temperature tolerance and the interdependent nature of these two traits in *S. cerevisiae*.

## INTRODUCTION

*Saccharomyces cerevisiae* is widely used to make fermented foods and beverages (Sicard and Legras 2011) and is the primary organism used to produce bioethanol (Azhar et al. 2017). Instrumental to *S. cerevisiae*’s utility is its preference for fermentation rather than respiration in the presence of oxygen (Merico et al. 2007) and its tolerance to high concentrations of ethanol (Stanley et al. 2010). Whereas the fermentative life-style evolved long ago and is shared by many other yeast species (Hagman et al. 2013), *S. cerevisiae*’s ethanol tolerance, a more recent acquisition, is shared only with other *Saccharomyces* species (Williams et al. 2015). Thus, there is considerable interest in identifying genes and genetic variation that contribute to *S. cerevisiae*’s high ethanol tolerance.

Ethanol tolerance in yeast is a complex phenotype, as it is influenced by available nutrients and growth substrates, as well as by environmental factors such as temperature and osmotic pressure (Casey and Ingledew 1986, d’Amore et al. 1990). The interaction between ethanol and temperature tolerance in yeast has been of particular interest due to the similarity of the ethanol and temperature stress response and also to observations that ethanol and temperature tolerance depend on one another (Casey and Ingledew 1986, Piper 1995). One mechanism that has emerged in mediating both ethanol and temperature tolerance is the plasma membrane composition and membrane fluidity. Membrane fluidity is influenced not only by phospholipid and sterol composition, but also by ethanol and temperature (Piper 1995, Ding et al. 2009). Furthermore, lipid composition has been found to play an important role in both ethanol and thermal tolerance (Thomas and Rose 1979, Suutari and Laakso 1994). Both ethanol and temperature tolerance have been central to applied investigations of ethanol production by yeast (Banat et al. 1998).

Various strategies have been used to understand the genetic basis of ethanol and thermal tolerance in yeast. Screens of the yeast deletion collection have uncovered several hundred genes involved in ethanol and temperature tolerance that cover a broad range of functional categories (Fujita et al. 2006; Van Voorst et al. 2006; Yoshikawa et al. 2009; Auesukaree et al. 2009; Ruiz-Roig et al. 2010; Nakamura et al. 2014). Genes have also been identified from experimental evolution with selection for high ethanol (Voordeckers et al. 2015) or temperature (Caspeta et al. 2014; Huang et al. 2018) tolerance. Finally, strains with different ethanol or temperature tolerance have been used to map quantitative trait loci (QTL) and the underlying genes responsible for these differences (Hu et al. 2007; Sinha et al. 2008; Yang et al. 2013). However, the majority of these genetic studies only examine ethanol or temperature tolerance, but not both.

In this study we undertook a genetic analysis of naturally occurring variation in resistance to ethanol at high temperature. We first identified a Chinese strain that was sensitive to ethanol at high temperature. Through crosses to two resistant strains, we identified two genes, *SEC24* and *PSD1* that are largely responsible for the ethanol and temperature sensitivity. Two amino acid substitutions in *SEC24* underlie the major quantitative trait locus (QTL) for ethanol tolerance at high temperature in both crosses. For temperature tolerance alone we found a single amino acid substitution in *PSD1* that underlies the major QTL in one of the crosses. Our results show that the genetic variation in tolerance to ethanol at high temperatures is largely distinct from either ethanol or temperature tolerance alone and is caused by a large effect mutation in *SEC24*, an essential gene that is a component of the COPII vesicle coat, required for cargo selection during vesicle formation in endoplasmic reticulum to Golgi transport (Miller et al. 2002).

## MATERIALS AND METHODS

### Yeast strains

The screen for variation in ethanol and temperature tolerance was carried out using 15 wild yeast strains from different regions of China (Wang et al. 2012), 15 diploid wild strains from North America (Sniegowski et al. 2002; Fay and Benavides 2005; Hyma and Fay 2013), and one strain each from the Philippines and Nigeria (Fay and Benavides 2005). Homozygous diploids of the strains from China were made for the initial phenotype assays. Forty-nine haploid strains from the Yeast Gene Deletion Collection (Winzeler et al. 1999) were used for non-complementation tests with HN6. All the strains used in this study are described in Table S1 and Table S4.

### Phenotype assays

Strains were grown on YPD (rich medium) agar plates (1% yeast extract, 2% peptone, 2% glucose, 2% agar) supplemented with 0% - 10% ethanol at 25°, 30°, 37° and 40°. Strains were pinned from microtiter plates to Singer Plus plates, using a Singer RoTor © HDA, to a density of 384 colonies per plate. Spot dilutions were made by diluting overnight cultures 1:50 with water and then making 1:10 serial dilutions.

### QTL mapping

Two sets of haploid recombinant strains were generated for QTL mapping using the ethanol and temperature sensitive strain HN6. A set of 58 recombinants was generated from a cross between HN6 and YJF153, and a set of 73 recombinants was generated from a cross between HN6 and SD1. For each cross haploid parental strains were mated and sporulated, and tetrads were dissected using a Singer System MSM 200 microscope. Recombinant strains were phenotyped by measuring colony size (mm) after 2 days of growth at 37° on YPD and YPD supplemented with 6% ethanol (Table S2).

Recombinant strains were genotyped using RAD-seq (restriction site associated sequencing, (Baxter et al. 2011) as previously described (Hyma and Fay 2013). DNA was extracted, digested with Mfe1 and Mbo1 and ligated to one of 48 barcoded sequencing adaptors (Table S3). Ligated products were pooled into groups of 48, size selected (150-500 bp) by gel extraction and amplified using extension primers containing one of three different indexes (Table S3). Amplified pools were quantified, pooled at equal concentrations and sequenced on an Illumina HiSeq2500. A total of 92 million reads was mapped to the S288c reference genome using BWA (v0.7.5a, Li and Durbin 2009), yielding a median of 600k mapped reads per strain. SNPs were called using the GATK unified genotyping algorithm (McKenna et al. 2010) and variants were filtered using vcftools (Danecek et al. 2011) to eliminate sites with >10% missing data, qualities less than 60 and more than two alleles. In total, 893 markers were segregating across the genome in both crosses (HN6xYJF153 and HN6xSD1) and were used for mapping (Data File S1).

R/qtl (Broman et al. 2003) was used for linkage analysis. The Haldane mapping function was used to estimate recombination rates, and phenotypes were mapped using the EM algorithm with a non-parametric phenotype model. Bayesian 95% confidence intervals were obtained for the quantitative trait loci (QTL) on chromosomes 9 and 14 using R/qtl.

### Candidate gene complementation tests

Genes within QTL intervals were tested for complementation using two approaches: MoBY plasmid rescue and hemizygosity tests of non-complementation using the yeast deletion collection (Table S4). MoBY ORFs, Molecular Barcoded Yeast plasmids (Ho et al. 2009), were transformed into a heat and ethanol sensitive recombinant strain (YJF2630) for genes within the chromosome 9 QTL and into a heat sensitive recombinant strain (YJF2702) for genes within the chromosome 14 QTL. Both recombinants were from the from the HN6 × YJF153 cross. For tests of non-complementation, HN6 was mated to BY4741 strains carrying single gene deletions within both the chromosome 9 and the chromosome 14 QTL regions. The colony area of HN6 × Knockout hybrids was determined using ImageJ. A t-test was applied to find knockouts that failed to complement the defect.

### Cloning

DNA from genes and their promoter regions was amplified by PCR and cloned into *Bam*HI/*Bgl*II cut, gel purified CEN plasmid pAG26, a gift from John McCusker (Addgene plasmid #35127) by gap repair in yeast. Hi copy plasmids were made by cloning Oak alleles of *SEC24* and *PSD1* into *Spe*I/*Xma*I cut, gel purified pRS42H (EUROSCARF #P30636, Taxis and Knop 2006) in a similar manner. Chimeric constructs were generated by PCR and gap repair of the desired parental segments. Primers used in this study are in Table S3. Allele specific restriction digests were used to confirm the constructs.

### Rho°mutants

0Strains were grown to saturation in the dark twice in minimal medium supplemented with 25ug/ml ethidium bromide at 30°to produce Rho° mutants, which were confirmed by streaking for single colonies on YPGly (1% yeast extract, 2% peptone, 5% glycerol, 2% agar).

### Site Directed Mutagenesis

PCR reactions were run using Pfu Ultra II Fusion HS DNA polymerase (Agilent Technologies, Santa Clara, CA), and pAG26 with inserts of either the YJF153 version of *SEC24* or the YJF153 version of *PSD1* as template. XL 10-Gold Ultracompetent Cells (Agilent Technologies, Santa Clara, CA) were transformed separately with 4 PCR reactions for each mutant, and DNA from the resulting transformed strains was sequenced.

Amino acid altering mutations were generated in the YJF153 allele of *SEC24*, and mutant DNA from four independent PCR reactions was transformed into HN6. Phenotypes of the resulting strains were tested on YPD with and without 6% ethanol at 30° and at 37°.Three constructs from each *SEC24* mutant were sequenced, and 7 of 12 were correct.

Amino acid altering mutations were generated in the YJF153 allele of *PSD1*, and mutant DNA from four independent PCR reactions was transformed into YJF2703 (very heat sensitive). Phenotypes of the resulting strains were tested on YPD at 30° and at 37°. Three constructs from each *PSD1* mutant were sequenced, and 8 of 9 were correct.

All strains and constructs are available upon request to the corresponding author.

## RESULTS

To identify naturally occurring variation in temperature and ethanol resistance we screened 25 diploid strains of diverse origin (Table S1) by pinning to rich medium plates supplemented with 0-10% ethanol and growing at 37°. Heat alone was sufficient to severely restrict the growth of the HN6 diploid and had a moderate effect on three other diploid strains: HN9, SX6 and HLJ2 (Figure 1). Although most strains were able to grow at 37° on plates supplemented with ethanol, three strains from primeval forests in China, HN6, HN9 and HN14, were more sensitive to ethanol at high temperature than the others. Several strains, including the China strain SD1, and especially the North American Oak strain YPS163, tolerated growth at 37° with concentrations of ethanol as high as 6% - 8%. None of the strains showed much variation in growth at 30o with or without 6% ethanol (Figure S1).

**Figure 1.**
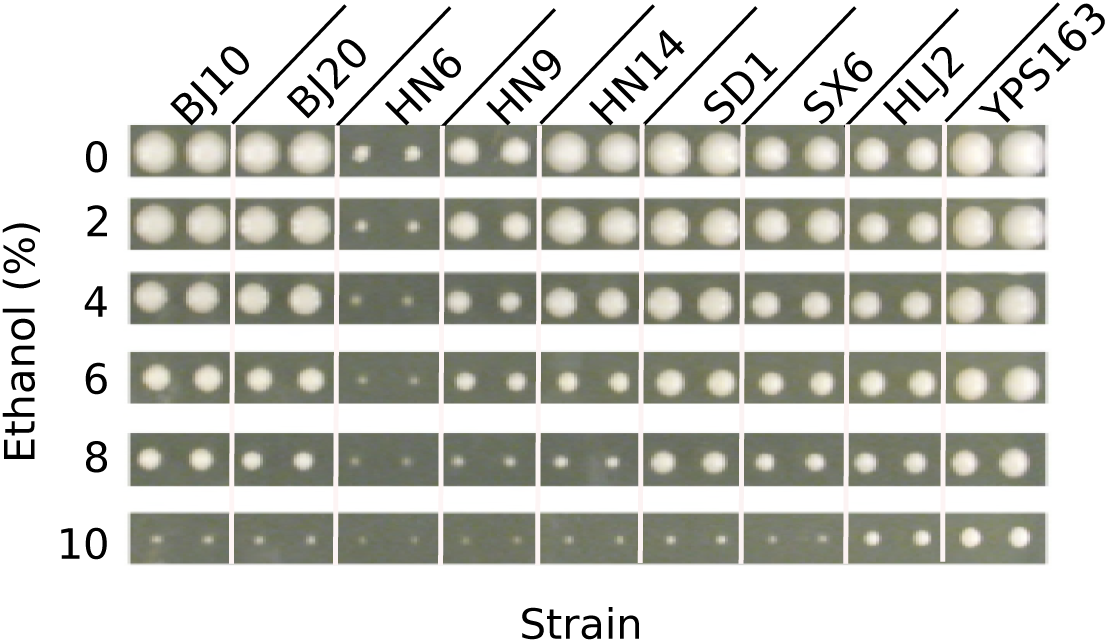
Ethanol and temperature dependent variation in colony size. Nine diploid strains were pinned in duplicate onto rich medium plates at different ethanol concentrations (0-10%) and grown at either 37° or 30° (Figure S1). Pictures were taken after 3 days of growth.

HN6, the most sensitive China strain and two heat and ethanol resistant strains, SD1 (China, persimmon) and YJF153 (haploid derived from YPS163, North American Oak), were selected for quantitative trait (QTL) mapping. Both HN6 × YJF153 and HN6 × SD1 diploid hybrids were temperature and ethanol resistant, indicating that HN6 carries recessive temperature and ethanol sensitive alleles. We generated 58 and 73 recombinant progeny from the HN6 × YJF153 and HN6 × SD1 crosses, respectively. The 131 recombinants were phenotyped at 37° on rich medium plates with and without 6% ethanol and were genotyped at 893 markers across the genome using RAD-sequencing (Methods). Controls were the haploid parental strains and the hybrids. In contrast to the very poor growth of the homozygous diploid HN6 at 37° in the initial screens (Figures 1 and S1), the HN6 haploid controls showed moderate growth (see Figure 5a).

**Figure 2.**
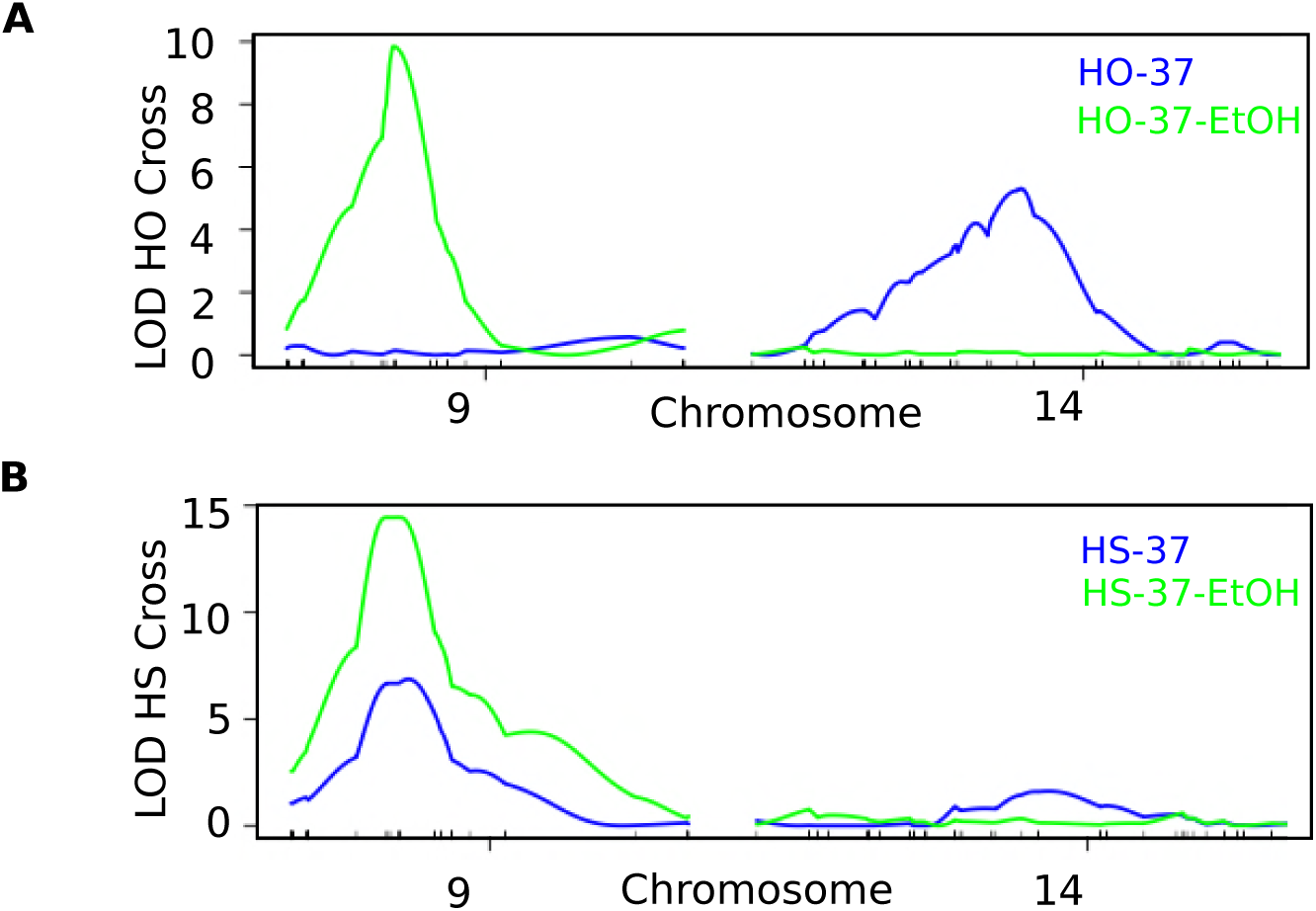
Quantitative trait loci for high temperature growth with and without ethanol. Logarithm of the odds ratio (LOD) of linkage across chromosomes 9 and 14, using (A) HN6 × YJF153 (HO) recombinants and (B) HN6 × SD1 (HS) recombinants, grown at 37° (blue) and 37° with 6% ethanol (green).

**Figure 3.**
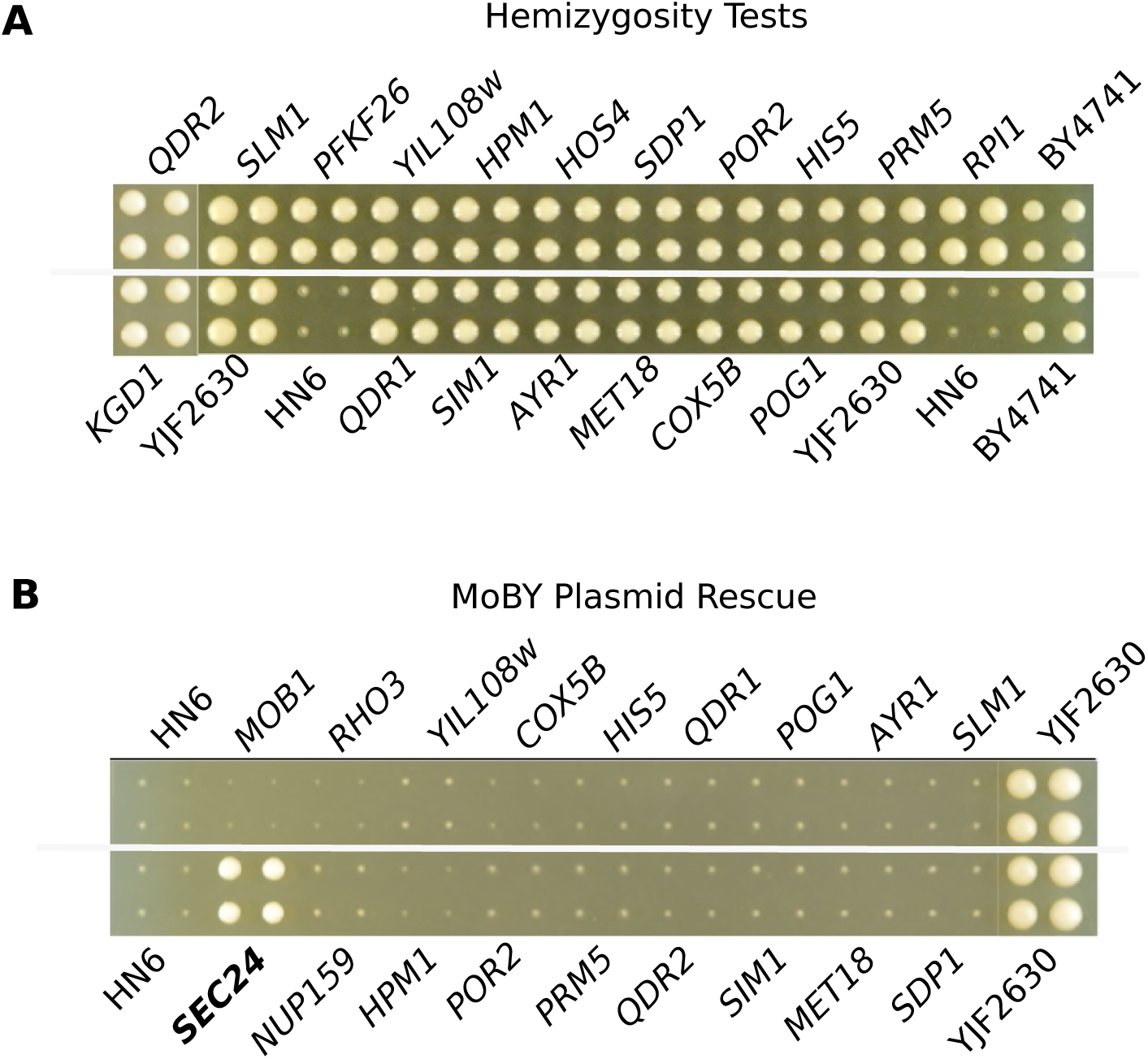
Complementation tests of genes within the chromosome 9 QTL identify *SEC24*. (A) HN6 × deletion strain hybrids grown at 37° with 6% ethanol. The control hybrid HN6 × BY4741 (YJF2630) and the parental strains HN6 and BY4741 are also shown. There are four replicate colonies for each strain with strain names above and below the colony images. All deletions complemented the no growth phenotype. (B) YJF2609 recombinant strains carrying MoBY plasmids were grown at 37° with 6% ethanol, four replicate colonies per strain. The HN6 and HN6 × BY4741 (YJF2630) controls are also shown. The SEC24 MoBY rescued the no growth phenotype of YJF2609.

**Figure 4.**
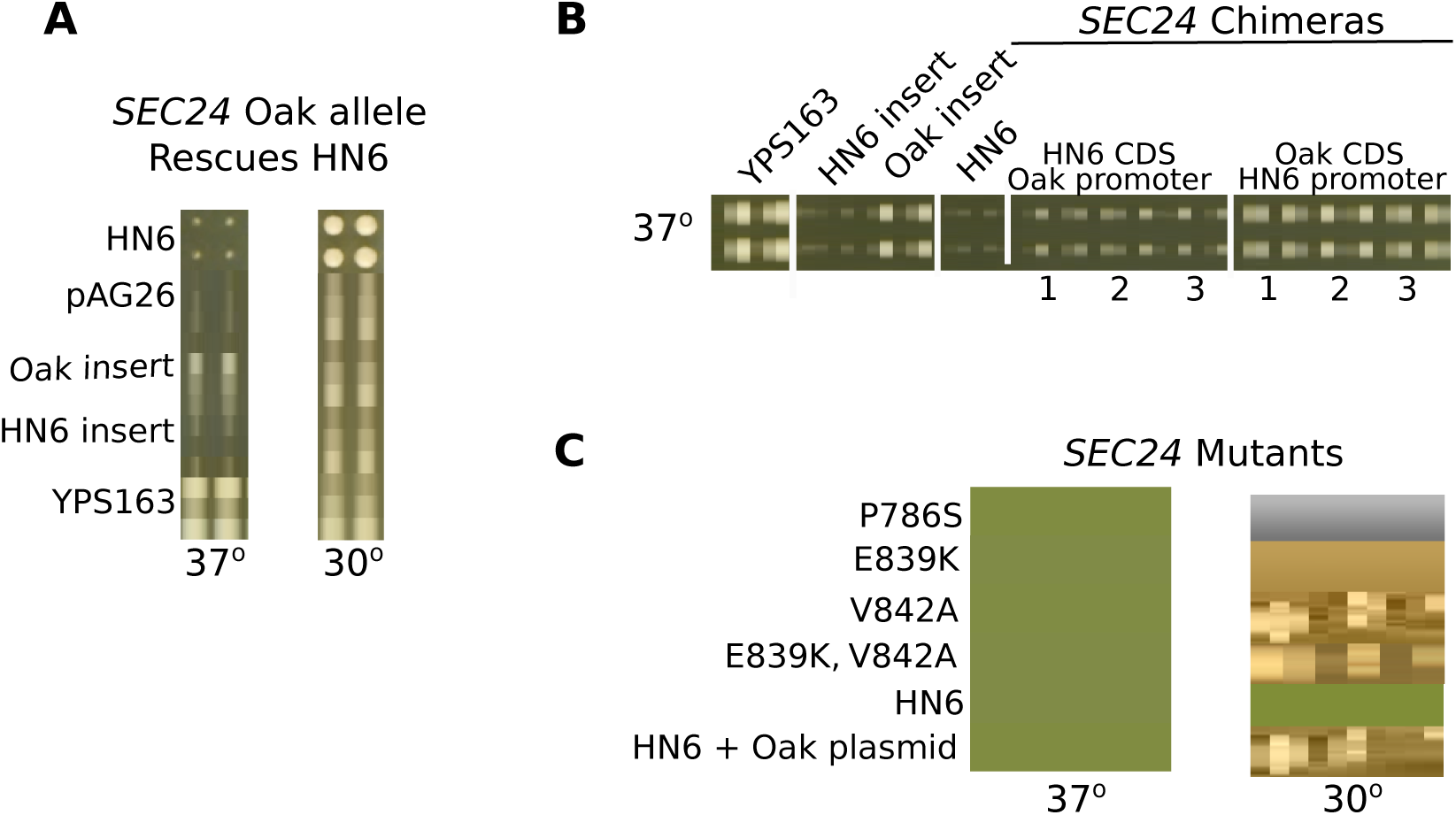
Two amino acid changes in *SEC24* are responsible for growth at 37° with ethanol. (A) Growth of HN6 at 37° with 6% ethanol is rescued by plasmids bearing the YJF153 allele of *SEC24* but not the HN6 allele of *SEC24* or the empty plasmid (pAG26). Growth at 30° with 6% ethanol is shown as a control. (B) Growth of HN6 at 37° with 6% ethanol is rescued by plasmids bearing the coding region of *SEC24* from YJF153. *SEC24* alleles with HN6 promoter and the YJF153 coding region rescue the phenotype, whereas alleles with the YJF153 promoter and HN6 coding region do not. Three transformants each are shown. The parental alleles for YJF153 and HN6 along with the YJF153 resistant and HN6 sensitive parental strains are controls. (C) Growth of HN6 at 37° with 6% ethanol is rescued by plasmids bearing mutant P786S, but not by mutants E839K or V842A, either as single mutants or as the double. The controls are the parent strain HN6 and YJF2696 (HN6 with the YJF153 plasmid).

**Figure 5.**
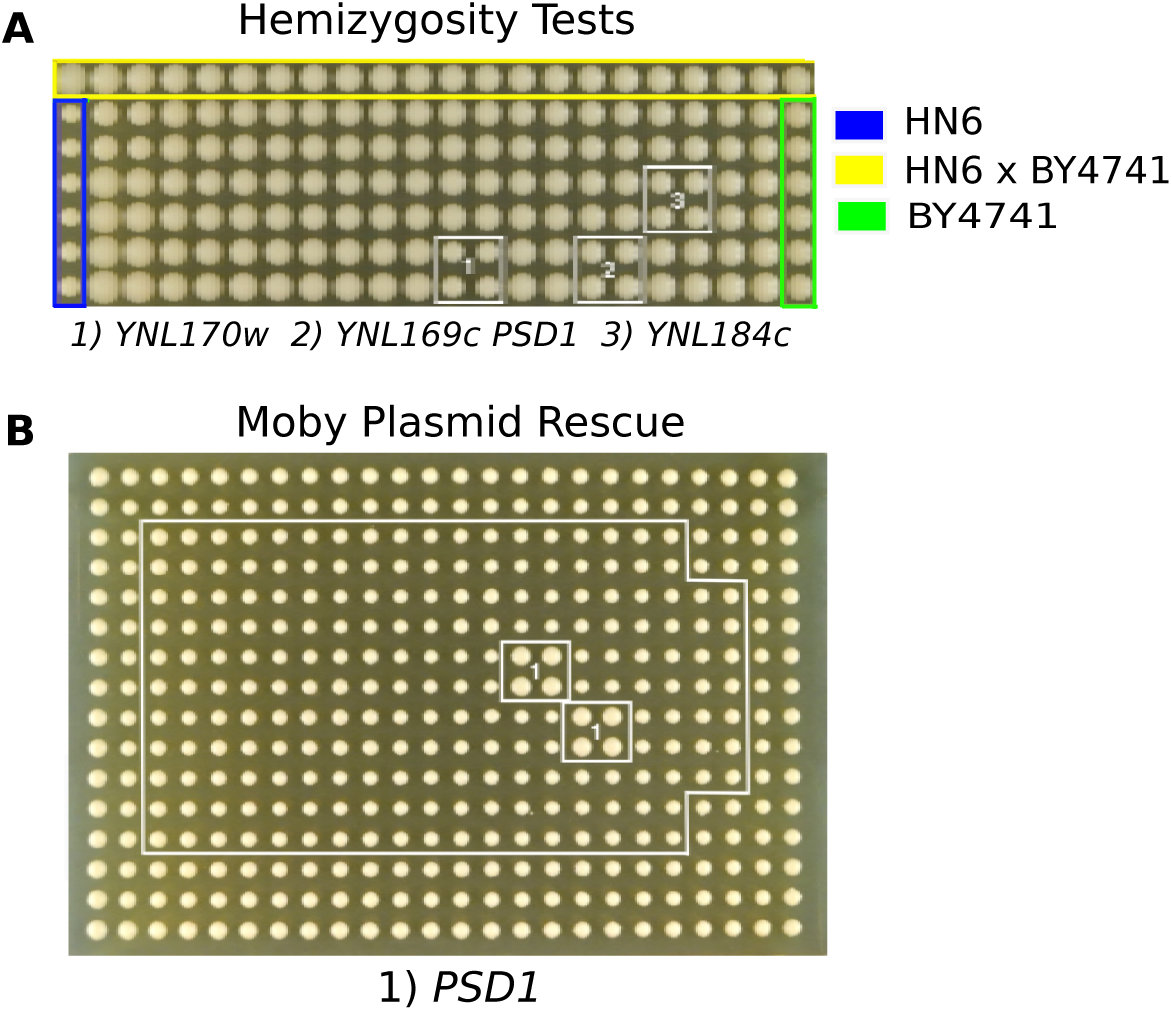
*PSD1* underlies the heat sensitivity on Chromosome 14. (A) Heat sensitivity of the HN6 × Knockout Hybrids, grown at 37° on YPD. The control hybrid YJF2630 (HN6 × BY4741), HN6 and BY4741 parental strains and three hemizygous strains that failed to complement the heat sensitive phenotype are labeled. Four replicate colonies are shown for each strain. (B) *PSD1* MoBY plasmid rescue of heat sensitivity of the recombinant strain YJF2702. YJF2702 without the plasmid is shown on the periphery (white lines), and the MoBY plasmid bearing strains are shown in the middle with four replicate colonies per strain. The strain bearing the *PSD1* plasmid is outlined by white boxes in two locations on the plate.

In both sets of recombinants approximately 50% of the strains were able to grow at 37° with 6% ethanol, indicating a single, major effect QTL. Consistent with this finding, we identified a single QTL in both crosses for growth at 37° with ethanol on chromosome 9 (Figure 2). The recombinant strains showed a more continuous distribution of growth at 37° without ethanol (Table S2). We identified a single QTL on chromosome 9 for the HN6 × SD1 recombinants, but a different QTL on chromosome 14 for the HN6 x YJF153 recombinants (Figure 2).

### Complementation tests

We used two different approaches to test genes within each QTL for their effects on temperature and ethanol. Because the diploid hybrids made with YJF153 are temperature and ethanol resistant, we would expect that (1) addition of a causal allele on a plasmid from a temperature and ethanol resistant strain into HN6 should increase growth, and that (2) the absence of a causal allele in a diploid hybrid should reduce growth. For plasmid rescue experiments we used the MoBY collection of plasmids containing the coding and adjacent noncoding regions of the lab strain S288c, which is both temperature and ethanol resistant. For the hemizygosity analysis we mated the sensitive strain, HN6, to BY4741 deletion strains, which are derivatives of S288c.

### Two amino acid substitutions in *SEC24* are responsible for the chromosome 9 QTL

There are 24 genes within the 56kb region on chromosome 9 associated with growth on ethanol at 37° (Table1). Single gene deletion strains were available for 18 of the genes in this region; MoBY plasmids were available for 18 of the genes, and neither deletion strains or plasmids were available for two genes (Table S4). None of the hybrids generated by crossing HN6 to the deletion strains were sensitive to growth at 37° with ethanol (Figure 3a). However, a single MoBY plasmid carrying *SEC24,* an essential gene, rescued growth of a recombinant strain (YJF2609) at high temperature with ethanol (Figure 3b). We used a recombinant strain that did not grow at 37° with ethanol rather than HN6, because the recombinant did not carry the G418 resistance gene (KAN), needed to select for the MoBY plasmids. Since *SEC24* is an essential gene in S288c, it could not be tested for non-complementation using deletion strains.

**Table 1.** Candidate genes tested for two QTL regions.

| QTL | Region (size) | ORFs | MoBY tested | Deletion tested | Not tested |
| --- | --- | --- | --- | --- | --- |
| Chr9 | 113,806-169,641 (~56 kb) | 24 | 18 | 18 | 2 |
| Chr14 | 258,375-323,565 (~65 kb) | 39 | 28 | 31 | 2 |

To confirm *SEC24* as a causal gene we tested the HN6 and YJF153 alleles of *SEC24* for plasmid rescue and found only the YJF153 allele rescued growth (Figure 4a). To identify causal variants within *SEC24* we first generated chimeric constructs between the HN6 and YJF153 alleles. Plasmids bearing the HN6 promoter region and the YJF153 coding region rescued growth at high temperature with ethanol, whereas those bearing the YJF153 promoter region and the HN6 coding region did not (Figure 4b).

To identify causal mutations within *SEC24* we tested three non-synonymous changes present in the coding region, all of which are near the 3’ end of the gene (Table 2). SIFT predictions (Sim et al. 2012) suggest that P786S and E839K affect gene function and that V842A will be tolerated. The Fungal Orthogroups Repository (Wapinski et al. 2007) shows high conservation for P786S, less for E839K and very little for V842A.

Phenotypic tests of the three non-synonymous changes in the YJF153 allele of *SEC24* grown with 6% ethanol at 37°, gave the following results: mutant P786S grew to the same extent as the positive control YJF2696; mutants E839K and V842A failed to grow. The double mutant E839K, V842A also failed to rescue growth (Figure 4c).

### A single amino acid substitution in *PSD1* is responsible for the chromosome 14 QTL

There are 39 genes within the 65kb region on chromosome 14 associated with growth at 37° without ethanol (Table1). Single gene deletions were available for 31 of the genes in this region; MoBY plasmids were available for 28, and only two genes were not represented by either a deletion strain or MoBY plasmid (Table S4). Three of the deletions (YNL170W, YNL169C/*PSD1* and YNL184C) showed effects in the hemizygous diploids (Figure 5a), and only one of the MoBY plasmids, bearing *PSD1*, rescued high temperature growth of YJF2702 (Figure 5b). The small ORF YNL170w, which had a phenotype in the hemizygosity test, has an overlap of 205bp with the 5’ end of *PSD1*, which includes G489R, the mutant responsible for the heat sensitive phenotype (see below). Plasmid rescue was conducted in the heat sensitive recombinant strain YJF2702 since it does not carry the KAN gene, required for maintenance of the MoBY plasmids.

To confirm that allele differences in *PSD1* affect high temperature growth, plasmids bearing the YJF153 and HN6 alleles of *PSD1* were tested for complementation. While the YJF153 allele of *PSD1* did not rescue growth of HN6 at 37° (Figure 6a), it was able to rescue high temperature growth of two recombinant strains, YJF2702 (mildly sensitive) and YJF2703 (very sensitive) (Figure 6b). Both of these recombinant strains grow at 37° on rich medium with 6% ethanol, and YJF2702 has a larger colony size (5mm) than YJF2703 (4mm). Both have the YJF153 genotype at *SEC24*.

**Figure 6.**
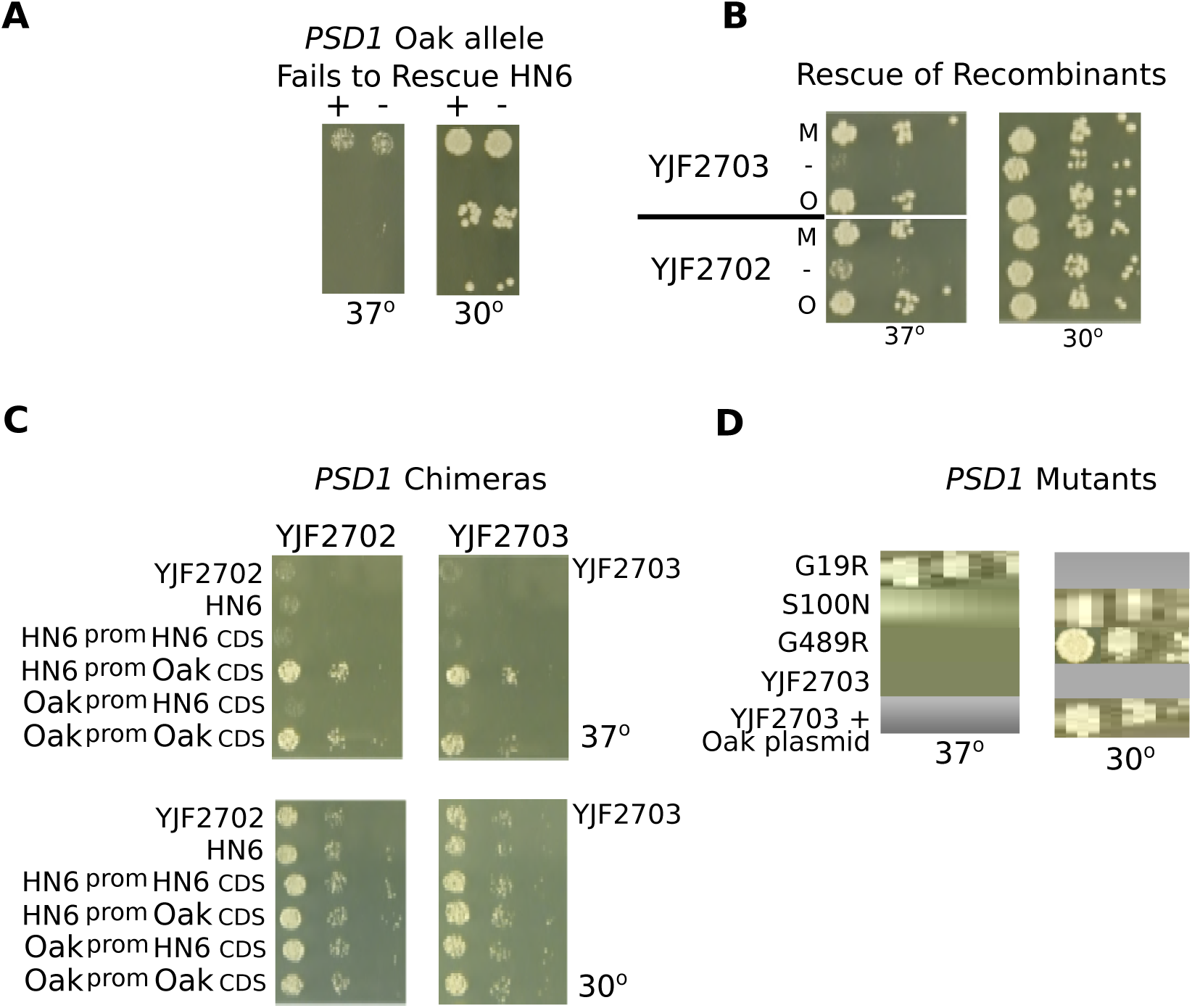
Rescue of the heat sensitive phenotype by YJF153 alleles of *PSD1*. (A) Spot dilutions show that the *PSD1* YJF153 plasmid fails to rescue the heat sensitive phenotype in the HN6 parent strain grown on YPD at 37°. (B) Recombinant strains YJF2702 and YJF2703 are rescued by the *PSD1* YJF153 plasmid (O) and by the *PSD1* MoBY (M) but not an empty vector (-). (C) YJF2702 and YJF2703 were transformed with chimeric *PSD1* plasmids, which contained either the YJF153 promoter joined to the HN6 CDS, or the HN6 promoter with an YJF153 CDS. The chimeric HN6 promoter YJF153 CDS plasmid rescues the phenotype to the same extent as the control YJF153 plasmid, indicating that the CDS is responsible for rescue. The YJF153 promoter HN6 CDS plasmid and the control HN6 plasmid fail to rescue. (D) Mutating the three non-synonymous sites in *PSD1* shows that mutant G489R fails to rescue YJF2703 on YPD at 37°. Mutants G19R and S100N rescue heat sensitivity to the same extent as the positive control, YJF2908 (YJF2703 with the YJF153 plasmid).

One potential difference between the recombinant strains and HN6 is their mitochondria; the recombinant strains could inherit either the HN6 or YJF153 mitochondria. Because *PSD1* functions in the mitochondria, and its deletion affects mitochondrial phenotypes, we tested whether (1) the heat sensitivity of HN6 depended on the mitochondrial type and (2) whether *PSD1* rescue depended on the mitochondrial type. To test for mitochondrial effects we generated HN6 × YJF153 hybrids with either HN6 mitochondrial DNA, YJF153 mitochondrial DNA or no mitochondrial DNA. Only hybrids lacking a mitochondrial genome showed reduced growth, as expected for a petite mutant (Figure 7a). To test whether rescue by the YJF153 allele of *PSD1* in the recombinant strains depends on mitochondrial type, we generated 2 recombinants carrying the YJF153 (Oak) allele of *PSD1* on a plasmid, but lacking mitochondrial DNA (Rho°). Both YJF2702 and YJF2703 that contained the plasmid but had no mitochondrial DNA, grew nearly as well as the strains containing both the plasmid and functioning mitochondria, showing that Rho° mutants of these strains can be rescued. YJF2702 that contained functional mitochondria but no plasmid, grew moderately well. The more heat sensitive recombinant YJF2703 behaved differently: in this case the strain contained functional mitochondria but no plasmid and grew nearly as poorly as the negative control (Figure 7b).

**Figure 7.**
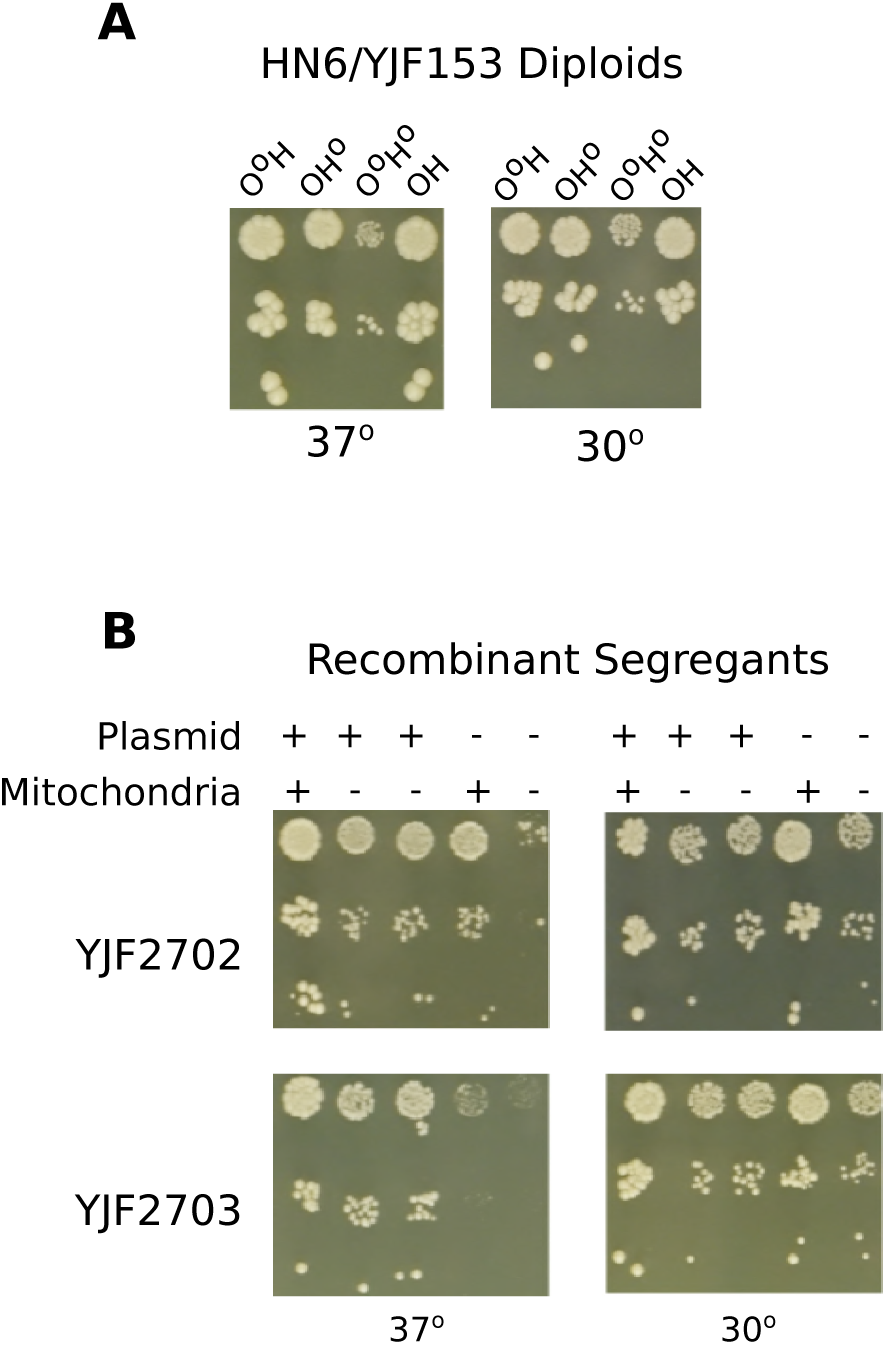
Mitochondrial involvement in the rescue of heat sensitive strains. (A) Spot dilutions of HN6/YJF153 diploids with mitochondrial DNA inherited from either parent, both parents, or no mitochondria show that strains are able to grow well at 37° with mitochondria from either parent. The O° H diploid lacks Oak mitochondria; H°O, lacks HN6 mitochondria; O°H° has no mitochondria; OH is derived from Rho^+^ parents. (B) Spot dilutions of heat sensitive recombinant strains YJF2702 and YJF2703 from the HN6 × YJF153 diploid (YJF2609) show that both YJF2702 and YJF2703 containing the plasmid, but had no mitochondria, grew nearly as well as the strains containing both the plasmid and mitochondria.

To identify causal variants in *PSD1*, we generated chimeric constructs between the HN6 and YJF153 alleles. Plasmids bearing the HN6 promoter region and the YJF153 coding region rescued growth of two recombinants, YJF2702 (mildly sensitive) and YJF2703 (very sensitive), at high temperature, whereas those bearing the YJF153 promoter region and the HN6 coding region did not (Figure 6c). We found three non-synonymous differences between the two alleles of *PSD1* (Table 2). According to SIFT predictions (Sim et al. 2012), only mutant G489R, quite close to the 3’ end of the gene, is likely to affect function; mutants G19R and S100N should be tolerated. The Fungal Orthogroups Repository (Wapinski et al 2007) showed extremely high conservation for mutant G489R and very little conservation in the other two.

**Table 2.** Non-synonymous changes in *SEC24* and *PSD1*.

| Gene | Allele |
| --- | --- |
| *SEC24* | P786S |
|  | E839K* |
|  | V842A* |
| *PSD1* | G19R |
|  | S100N |
|  | G489R* |

Phenotypic tests of the three non-synonymous changes in the YJF153 allele of *PSD1* grown at 37°, gave the following results: mutant G19R and S100N rescued the heat phenotype, whereas mutant G489R failed to rescue (Figure 6d).

### Overexpression of the Oak alleles of *SEC24* and *PSD1* does not enhance tolerance to heat and ethanol or to heat alone in sensitive strains or in wild strains of yeast

To test whether overexpression of the Oak alleles of *SEC24* and *PSD1* would further enhance tolerance of sensitive strains to heat and ethanol or to heat alone, we transformed them with hi copy plasmids bearing these alleles. HN6 bearing the Oak allele of *SEC24* in high copy grew very poorly on YPD at 37° compared with the same strain with the gene in a CEN plasmid. The CEN plasmid version of the strain showed somewhat less growth on YPD+ethanol at 37°; however the strain with the high copy plasmid did not grow at all. (Figure 8a). Although rescued by the Oak *PSD1* CEN plasmid, the heat sensitive strain YJF2703 bearing an overexpressed Oak allele of *PSD1* grew poorly on YPD at 37° (Figure 8b).

**Figure 8.**
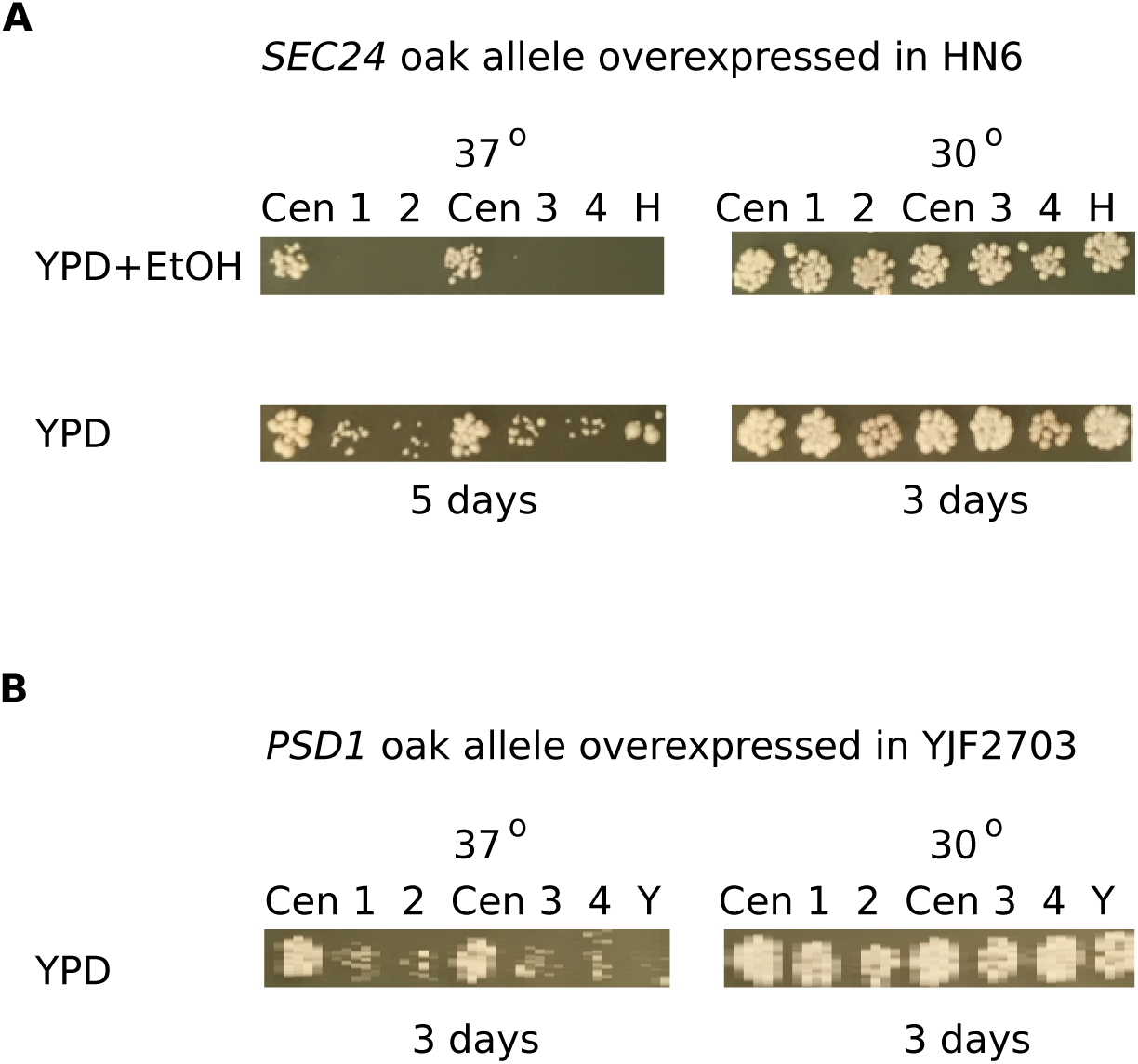
Phenotypes of sensitive yeast strains containing high copy Oak allele plasmids grown at high temperature on YPD+ethanol and on YPD. (A) HN6 is rescued on YPD+ethanol at 37° by the *SEC24* CEN plasmid, but the high copy version is lethal. On YPD alone at high temperature HN6 with the high copy plasmid grows very poorly (5 days of growth). The 30° controls are shown at 3 days of growth. There are four transformants (1 - 4) of each high copy plasmid. H = HN6, Cen = CEN plasmid. (B) YJF2703 is rescued on YPD at 37° by the *PSD1* CEN plasmid, but the high copy version is deleterious. These and the 30° controls are shown at 3 days of growth. There are four transformants (1 - 4) of each high copy plasmid. Y = YJF2703, Cen = CEN plasmid.

The effects of *SEC24* and *PSD1* on resistance to heat and ethanol and resistance to heat alone raise the possibility that increasing the activity of these genes might enhance heat and ethanol resistance in strains that carry the resistant *SEC24* and *PSD1* alleles: Oak, Wine and S288c. We tested whether overexpression of *SEC24* and *PSD1* would improve resistance using high copy plasmids. Since 37° is a permissive temperature for the resistant Oak, Wine and S288c strains, 40° was used for phenotype testing.

The presence of high copy or CEN plasmids bearing the Oak alleles of *SEC24* or *PSD1* did not enhance the growth of the Oak, Wine or S288c strains grown on YPD at 40°; and there was no difference in growth between the CEN and high copy plasmids in these strains (Figure S2a). Although growth was abundant on YPD at 40° by day 3, there was little or no growth on YPD+ethanol at 40° (Figure S2b).

## DISCUSSION

*S. cerevisiae* is known for its resistance to high concentrations of ethanol (Arroyo-López et al. 2010). However, ethanol resistance is also known to be temperature dependent (Casey and Ingledew 1986). Prior studies have either focused on genetic variation in either ethanol or temperature stress but not both (Khatun et al. 2017; Kitichantaropas et al. 2016; Nuanpeng et al. 2016; Benjaphokee et al. 2012). In this study we mapped ethanol tolerance at high temperature and identified two large effect amino acid substitutions in *SEC24*. We also identified a smaller effect amino acid substitution in *PSD1* that caused resistance to heat alone. Our results provide new evidence for the involvement of ER to Golgi transport in ethanol tolerance at high temperature (*SEC24*) and support prior work on the role of mitochondrial fusion in temperature tolerance (*PSD1*).

Differences in genetic background lead us to identify one gene involved in heat or heat and ethanol tolerance (*SEC24*) and one gene involved in heat tolerance alone (*PSD1*) through quantitative trait mapping in two crosses. Both crosses shared the parental strain HN6, a heat and ethanol sensitive strain isolated from a primeval forest in Hainan, China (Wang et al. 2012). However, the two crosses differed in the other parent (YJF153 or SD1) and the effects of the sensitive *SEC24* and *PSD1* alleles from HN6. In the HN6 × SD1 cross, the HN6 *SEC24* allele conferred sensitivity to heat and even greater sensitivity to heat and ethanol combined. However, in the HN6 × YJF1533 (Oak) cross, the HN6 *SEC24* allele only caused sensitivity to heat and ethanol combined whereas the HN6 allele of *PSD1* caused sensitivity to heat alone. Interestingly, the sensitivity of HN6 was not as severe in the haploid compared to the diploid version, which might have been caused by a dosage effect. Both HN6 defects could be rescued by plasmids carrying the corresponding alleles of *PSD1* and *SEC24* from YPS163, a resistant strain collected from an oak tree in Pennsylvania, USA (Sniegowski et al. 2002). However, in the case of *PSD1*, rescue only occurred in HN6 × YJF153 recombinant strains bearing the *PSD1* sensitive allele, indicating that alleles from the YJF153 background are necessary for expression of *PSD1* heat tolerance. We refined the effects of the HN6 alleles to a single amino acid substitution in *PSD1* and either of two substitutions in *SEC24*.

The phenotypic effects of amino acid substitutions in *SEC24* point to its importance in the combined tolerance to ethanol and high temperature. However, because *SEC24* is an essential gene it is also possible that the amino acid substitutions result in temperature sensitive alleles of *SEC24*, and that *SEC24* is not inherently involved in ethanol and heat tolerance. Our observation that *SEC24* alleles do not influence sensitivity to heat alone in one of the two crosses (HN6 × YJF153) does not support this later interpretation. Another potential mechanism of *SEC24* mediated sensitivity to ethanol and heat is its role in ER to Golgi transport. ER to Golgi transport is an important component of protein quality control; misfolded proteins in the ER are transported back into the cytoplasm in order to be degraded by the ubiquitin–proteasome system (Taxis et al. 2002). While it is not obvious why the *SEC24* allele from HN6 is particularly sensitive to heat in the presence of ethanol, this phenotype may be mediated by defects in the transport of proteins important to heat and ethanol tolerance or to defects in the Golgi or ER membranes themselves.

The mechanism by which *PSD1* affects heat sensitivity is likely related to its impact on mitochondrial membranes, but depends on other genetic factors coming from the YJF153 background. *PSD1* converts phosphatidylserine to phosphatidylethanolamine (PE), a mitochondrial phospholipid that plays an important role in mitochondrial fusion and in the maintenance of mitochondrial morphology (Joshi et al. 2012, Zhang et al. 2014). Mitochondrial function is known to be important for intrinsic heat resistance, and deletion of two genes, *CHO1* and *OPI3*, required for conversion of PE to phosphatidylcholine results in heat shock sensitivity (Jarolim et al. 2013). Furthermore, it has been proposed that heat induced changes in membrane fluidity influence the perception of high temperature and the expression of heat shock proteins (Carratù et al 1996). While *PSD1* has not previously been identified as a gene conferring resistance to high temperatures, this may be a consequence of its dependence on unknown genetic factors segregating in the HN6 × YJF153 recombinants. Our complementation analysis indicates that this unknown factor is not the mitochondrial DNA type.

Overexpression of *PSD1* and *SEC24* did not enhance heat or heat and ethanol tolerance, and in some instances was toxic. The lethality in sensitive strains and failure to enhance growth in resistant strains could be caused by the fact that Sec24 is one of five essential proteins that form the COPII vesicle coat. It along with Sec23 forms the inner coat of the vesicle as a heterodimer and binds the cargo that will be transported from the rough Endoplasmic Reticulum to the Golgi apparatus. (Jensen and Schekman 2011 review). High copy expression of *SEC24* might lead to an overabundance of the protein, which in turn might hinder heterodimer formation. It has been shown that overexpression of both *SEC24* and *SEC23* leads to decreased growth (Sopko et al. 2006) and decreased growth rate (Yoshikawa et al. 2011) in yeast.

High copy expression of *PSD1* was harmful to heat sensitive strains and did not enhance the growth of wild type thermotolerant strains. These results are in line with previous studies showing that overexpression decreases the vegetative growth rate (Yoshikawa et al. 2011) and induces invasive growth (Shively et al. 2013). Overexpression of *PSD1* might hinder growth in yeast grown at high temperatures, as seen in our sensitive strains, since PE is normally only expressed at high levels during growth at low temperatures (Henderson et al 2013). Similar reasoning might explain the failure of *PSD1* to enhance growth in resistant strains grown at high temperature.

In summary, the HN6 alleles of SEC24 and PSD1 explain its sensitivity to heat and ethanol. However, the effects of both genes differ depending on the cross, implying that there are other differences between the YJF153 and SD1 background that modify their effects. We conclude that ethanol and temperature tolerance are worth examining together in studies of quantitative genetic variation in S. cerevisiae.

## Acknowledgements

We thank members of the Fay lab for their comments and suggestions. This work was supported by an NIH grant (GM080669) to J. Fay.

---

**Supplemental Table 1.** Strains used in this study.

**Supplemental Table 2.** Recombinant strain barcode indices, sequencing and phenotypes used for QTL mapping.

**Supplemental Table 3**. Primers used in this study.

**Supplemental Table 4.** Genes tested for complementation using a MoBY and/or a hemizygosity test.

**Supplementary Figure 1**. Temperature and ethanol sensitivity of twenty-five *S. cerevisiae* strains. Strains were grown in quadruplicate at either 30° or 37° on rich medium (YPD) without ethanol or with 6% ethanol. The layout of the strains is shown by the names below the images and the strains chosen for further study (HN6, SD1 and YPS163) are in bold and are outlined in white.

**Supplementary Figure 2.** Phenotypes of resistant wild yeast strains, Oak, Wine and S288c, containing high copy Oak allele plasmids grown at 40°. (A) Growth on YPD at 40° is diminished for Wine and S288c, but not for Oak. (B) High copy plasmids are deleterious for all three wild strains grown at 40° on YPD+ethanol. The controls were grown at 37°.All are shown at three days of growth. The YPD grown strains are shown at a 10-fold higher dilution than those grown on YPD/EtOH. P* = *PSD1* high copy, P = *PSD1* CEN plasmid; S* = *SEC24* high copy, S = *SEC24* CEN plasmid; Par = parent strain.

**Data File S1.** Genotypes and phenotypes of recombinant strains used for QTL mapping.

